# Novel lytic and lysogenic cyanophages predicted to infect *Microcoleus* associated with anatoxin-producing benthic mats

**DOI:** 10.1101/2023.04.12.536658

**Authors:** Cecilio Valadez-Cano, Adrian Reyes-Prieto, Janice Lawrence

## Abstract

Proliferations of toxic benthic cyanobacteria are increasingly being reported around the world. Of particular concern are *Microcoleus*-dominated mats associated with anatoxin production that have resulted in dog fatalities. Although the impact of cyanophages has been demonstrated in planktonic systems, their role in the population dynamics of benthic cyanobacteria has received little attention. Here we use metagenomics to explore phage presence in benthic mats from the Wolastoq|Saint John River (WR; New Brunswick, Canada) and Eel River (ER; California, US). Our survey recovered multiple viral-like sequences associated with different putative bacterial hosts, including two cyanophage genomes with apparently different replication strategies. A lysogenic cyanophage (predicted as a prophage) was found integrated in the genomes of *Microcoleus* sp. 3 recovered from five ER mat samples. This Microcoleus phage is related to previously described Phormidium phage counterparts. Also, we recovered lytic cyanophages from WR and ER mats dominated by anatoxin-producing *Microcoleus*, which was predicted as the putative host. Despite the geographical distance between WR and ER, the lytic Microcoleus phage genomes recovered from each river have similar sizes (circa 239 Kbp) and share similar gene content with high sequence identity. Phylogenetic analysis suggests that these lytic Microcoleus phages are distant from any other cyanophage previously described. Our results constitute the first report of cyanophages predicted to infect and therefore influence the population dynamics of mat-forming *Microcoleus* spp. associated with anatoxin production.

## Introduction

Some species of benthic cyanobacteria (BC) have been associated with animal poisonings due to their potential to produce secondary metabolites with toxic effects (Wood et al., 2020). Among the toxins produced by BC are those with hepatotoxic (microcystins, nodularins and cylindrospermopsins), dermatoxic (lyngbyatoxin) and neurotoxic (saxitoxins and anatoxins) activities (Quiblier et al., 2013). Proliferations of toxigenic BC have been reported in lakes, reservoirs, streams, rivers, meltwater, and geothermal ponds in 19 countries (Wood et al., 2020). Toxigenic potential has been mainly described in cyanobacteria of the orders Nostocales (*Anabaena* and *Nostoc*) and Oscillatoriales (*Oscillatoria*, *Phormidium*, *Microcoleus* and *Microseira* genera) (Wood et al., 2020).

Numerous dog poisoning events in rivers have been associated with the exposure to anatoxins produced by mat-forming *Microcoleus* species in New Zealand (e.g., Wood et al., 2007), USA (California; e.g., Bouma-Gregson et al., 2018), and Canada (New Brunswick; e.g., McCarron et al., 2023). The high spatial and temporal variability in anatoxin concentration observed in benthic mats has been related with fluctuations in the abundance of toxigenic genotypes, rather than differential production per cell (Wood and Puddick, 2017). It is well documented that toxin-producing cyanobacteria species (e.g., genera *Microcoleus, Phormidium*), can co-occur with non-toxigenic congeneric relatives over small spatial scales (<1 cm) (Wood et al., 2012; Wood and Puddick, 2017).

Bacterial viruses (i.e., bacteriophages or phages) play important roles by shaping both cell abundance and diversity in microbial communities (Bouvier and Del Giorgio, 2007; Brown et al., 2019). In particular, phages infecting cyanobacteria (cyanophages) can impact the abundance and diversity of toxic cyanobacterial blooms (Grasso et al., 2022; Zhang et al., 2022). For example, the activity of cyanophages infecting microcystin-producing *Microcystis* species (Yoshida et al., 2008) can lead to a significant reduction in microcystin production (Naknaen et al., 2021). Reports of cyanophages infecting diverse planktonic bloom-forming cyanobacteria are abundant (see the review of Grasso et al., 2022), but in contrast the information about cyanophages associated with BC is limited. Few reports include cyanophages identified in mats from lakes (Voorhies et al., 2016) and rivers (Al-Shayeb et al., 2020), but the existence of phages associated with toxigenic mat-forming *Microcoleus* in riverine ecosystems remains unexplored.

While the number of cyanophage-cyanobacteria relationships explored to date is limited relative to the enormous estimated diversity of cyanophages, the use of high-throughput DNA/RNA sequencing technologies has led to unprecedented rates of virus discoveries directly from environmental samples (Al-Shayeb et al., 2020; Edwards et al., 2015; Nayfach et al., 2021b). For example, the use of these “shotgun sequencing” approaches has revealed viral genomic sequences from ancient ice samples (355-4,400 year old), predicted to infect co-occurring bacteria (Zhong et al., 2021). Also, thousands of complete phage genomes have been identified in human gut metagenomes infecting bacteria of the phylum Bacteroidetes (Benler et al., 2021).

Here we used high-throughput sequencing (Illumina) of DNA extracted from benthic mats collected along the Wolastoq (also known as Saint John River; New Brunswick, Canada) to both confirm the presence of anatoxin-producing *Microcoleus* and to explore the presence and diversity of phages. In parallel, we also investigated the presence of cyanophages in publicly-available Illumina sequences produced previously (Bouma-Gregson et al., 2019) from benthic mats from the Eel River (California, USA). Our survey of the metagenomic data obtained from both rivers revealed the presence of cyanophages likely infecting the *Microcoleus* species that dominate the benthic mats from these two geographically distant freshwater systems.

## Methods

### Sample collection

Samples of cyanobacteria-dominated benthic mats were collected during the summer-fall period of 2019 (Table 1) from 8 different sites along the Wolastoq (New Brunswick, Canada; Figure S1). Mat samples (approximately 5 g) were homogenized in 15 mL disposable tissue grinders (VWR International, Radnor, PA, USA). Once ground, homogenized samples were stored at −20 °C prior to extraction using the DNeasy^®^ PowerSoil^®^ Kit (Qiagen) following manufacturer’s instructions.

**Table 1.**
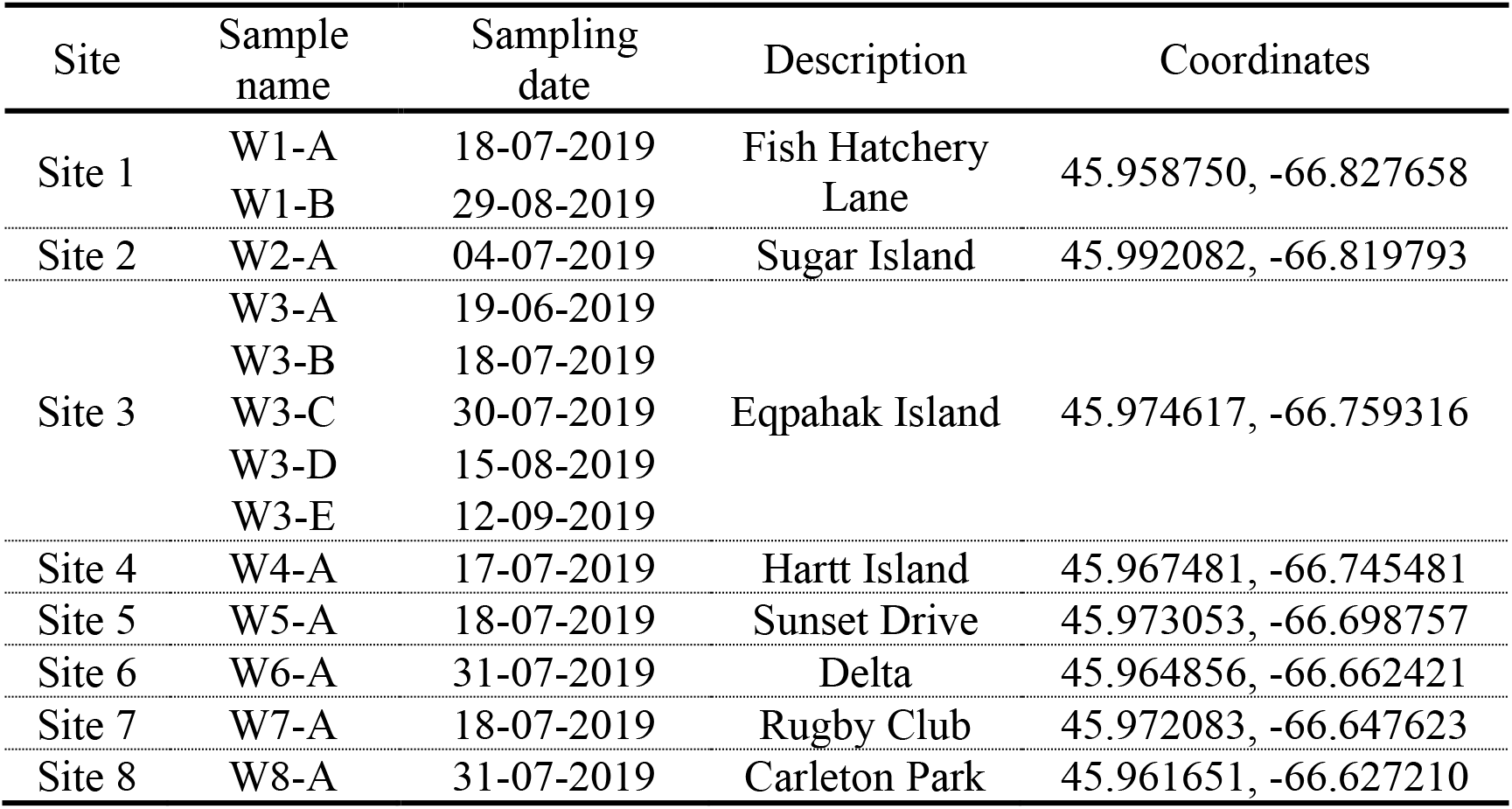
Site, name, sampling dates and location for benthic mat samples collected from the Wolastoq in 2019.

### Shotgun DNA sequencing

Total purified DNA from 13 mat samples was sequenced individually using Illumina technology (NextSeq 550 platform;150 bp paired-end reads). Preparation of libraries and sequencing were performed at the Integrated Microbiology Resource of of Dalhousie University (Halifax, NS) following standard protocols (https://imr.bio/protocols.html).

### Read processing and assembly

Illumina adapter removal and read trimming for quality control was performed with *fastp v0.20.1* (Chen et al., 2018). High quality reads were assembled with *metaSPAdes* (*SPAdes v3.12.0*) (Bankevich et al., 2012) using the k-*mer* options *-k21,33,55,77*.

In parallel, we retrieved from the NCBI Sequence Read Archive (SRA) raw Illumina sequencing sets of 61 Oscillatoriales-dominated microbial mats collected in 2015 (22 sets) and 2017 (39 sets) across the Eel River network (Northern California, USA; Bouma-Gregson et al., 2022, 2019). Quality control and assembly of the Illumina data sets was performed as described.

### Phage identification and lifecycle prediction

We used the *What the Phage* tool (*WtP v22.04.0*) (Marquet et al., 2022) to identify putative phage sequences in scaffolds larger than 20 Kbp from Wolastoq samples collected in 2019 and from the Eel River in 2015. Briefly, the *WtP* suite combines the scores of 12 tools designed to predict putative viral sequences in assembled contigs (Marquet et al., 2022). The quality of each predicted viral sequence was assessed with the *CheckV* tool (Nayfach et al., 2021a). Based on the results obtained from the *WtP* pipeline, candidate scaffolds were manually inspected to identify putative viral genomes. We predicted the replication strategy of the viral sequences with *Replicadec 0.2.3.1* (Peng et al., 2022). Briefly, *Replicadec* predicts the lifecycle of the predicted virus detecting well-described phage proteins (i.e., integrase and excisionase proteins) and genes (PI-like genes) associated with lysogenic and chronic replication strategies, respectively. *Replicadec*, also uses a Naïve Bayes classifier to calculate the conditional probability of the viral lifecycle based on alignment of the query viral genes with an in-house viral database.

### Phage-host prediction

We used information from both the CRISPR-Cas (Clustered Regularly Interspaced Short Palindromic Repeats) system and phage tRNA matches (Paez-Espino et al., 2016) in the metagenomic sequences to predict/identify putative bacterial phage hosts. In the case of the CRISPR-Cas system, we used the *CrisprOpenDB* database (comprises 11,767,782 spacer sequences from bacterial genomes) as a comparative reference (Dion et al., 2021). Also, we used the *Mining CRISPRs in Environmental Datasets* tool (*MinCED v0.4.2*) (Bland et al., 2007) to retrieve 15,495 spacer sequences from 178 MAGs obtained from *Microcoleus*-dominated mats and non-axenic isolates from different rivers in USA, New Zealand and Canada (Table S1) (Bouma-Gregson et al., 2022, 2019; Conklin et al., 2020; Tee et al., 2021; Valadez-Cano et al., 2023). All spacer sequences were compared against the putative phage sequences recovered from the Wolastoq and Eel River mats using the BLASTn-short function (E-value ≤ 1.0 x 10^-10^, sequence identity ≥ 95%, covering at least 90% of the query spacer sequence) of the BLAST+ package.

For host assignment based on tRNA matches, we used the *ARAGORN v1.2.38* tool (Laslett and Canback, 2004) to recover tRNA sequences from the 178 MAGs mentioned above. We also retrieved 1,019,860 bacterial tRNA sequences from the *tRNA gene database curated manually by experts* (*tRNADB-CE v13.0*) (Abe et al., 2009). The complete set of tRNA sequences were compared against the phage scaffolds from the Wolastoq and Eel rivers using again the BLASTn-short function of the BLAST+ package considering the 100% of both the query length and the sequence identity.

### Phylogenetic analysis

The phylogenetic affiliation of our predicted Microcoleus phages with cyanophages reported in the *Virus-Host DB* (Mihara et al., 2016) was evaluated using the genome-wide sequence similarity approach implemented in *ViPTree v3.0* (Nishimura et al., 2017). The *ViPTree v3.0* server utilizes the “Phage Proteomic Tree” concept developed by Rohwer and Edwards (Rohwer and Edwards, 2002) with some modifications (Nishimura et al., 2017). Briefly, *ViPTree* uses TBLASTX (BLOSUM45 substitution matrix) to compare all predicted proteins (the “proteome”) from the query genome against the reference database of viral genomes *GenomeNet/Virus-Host DB* (Mihara et al., 2016) to calculate normalized (correcting for differences in viral genome sizes) pairwise similarity scores. Then, the corrected TBLASTX scores are used to estimate genomic distances (*S_G;_* values from 0 to 1) between all compared viral sequences. Finally, the resulting dissimilarity matrix (1-*S_G_*) is used to construct a distance tree with BIONJ (Gascuel, 1997).

### Classification of recovered phage sequences

Taxonomic assignation of identified viruses to the order and family levels were performed with *Demovir* (Liao et al., 2022). First, the *Prodigal v2.6.3* (Hyatt et al., 2010) tool was used to predict Open Reading Frames (ORFs) in the putative viral genomes. Then, we used *Demovir* with default parameters to search for homologous proteins encoded in the genomic query and the viral subset of the TrEMBL database (Boeckmann et al., 2003). Taxonomic classification of Microcoleus phages at the genus level was performed with *vConTACT v0.9.22* (Bin Jang et al., 2019). Briefly, the protein sequences coded by the predicted Microcoleus phages and those coded by known viral genomes from the NCBI ViralRefSeq were grouped in protein clusters (PCs) using BLASTP. Groups of similar sequences were then clustered using the Markov clustering algorithm (MCL) with an inflation value of 2 to create viral clusters (VCs). The information identified from organized VCs and PCs were used to create module profiles, which can then be classified for taxonomic identification (Bin Jang et al., 2019).

To discard the possibility that predicted Microcoleus phages were plasmids, we used the *PlasFlow* pipeline (Krawczyk et al., 2018) available in the Galaxy Platform (https://galaxyproject.org/) (The Galaxy Community, 2022). *PlasFlow* uses genome signatures of sequences from different bacterial chromosomes and plasmids to train a deep neural network model to discriminate chromosomal and plasmid sequences from different phyla. Circularity of the Microcoleus phage genomes was explored with *find_circular.py* script implemented in the *ViRal and Circular content from metAgenomes* (*VRCA*) pipeline (Crits-Christoph et al., 2016)

### Annotation of predicted phage sequences

The sequences of the predicted Microcoleus phage genomes were annotated with the *Distilled and Refined Annotation of Metabolism v1.3* (*DRAM-v*) pipeline using default parameters (Shaffer et al., 2020). The *DRAM-v* pipeline uses the *MMSeqs2* (Steinegger and Söding, 2017) and *HHMER3* (Finn et al., 2011) tools to annotate the genes predicted with the *Prodigal v2.6.3* considering information from the Kyoto Encyclopedia of Genes and Genomes (KEGG; Kanehisa et al., 2017), Pfam (El-Gebali et al., 2019), dbCAN (Zhang et al., 2018), RefSeq viral (https://www.ncbi.nlm.nih.gov/genome/viruses/) and Virus Orthologous Groups database (VOGDB; https://vogdb.org/) databases.

Protein sequences predicted in the lytic Microcoleus phages were grouped into clusters of orthologs with *GET_HOMOLOGUES v3.3.2* (Contreras-Moreira and Vinuesa, 2013) using the orthoMCL algorithm (75% minimum coverage in BLAST pairwise alignments and max E-value 1e-05) (Li et al., 2003).

### Cyanobacterial relative abundance

The relative abundance of major cyanobacterial groups and anatoxin-producing *Microcoleus* species in the 2019 Wolastoq mats was estimated using a previously published approach (Valadez-Cano et al., 2023). Briefly, we recovered and annotated all detected *rps3* genes (documented to be single-copy in bacteria) in mat samples and assigned taxonomic affiliations to recovered sequences by similarity searches. Then, we mapped reads from each mat sample to non-redundant bacterial *rps3* genes and used the read coverage data (reads/nucleotide) to calculate the corresponding relative abundances in each mat sample.

To estimate the relative abundance of previously reported toxigenic and non-toxigenic *Microcoleus* subspecies from the Wolastoq (Valadez-Cano et al., 2023) in each sample, we used the corresponding read coverage of the *non-ribosomal peptide synthetase* cluster and the *anatoxin-a* cluster, which are exclusive to the non-toxigenic and toxigenic *Microcoleus* subspecies, respectively (Valadez-Cano et al., 2023).

### Sequencing coverage calculation of reads mapping to lytic Microcoleus phage genomes

We used *CoverM v0.6.1* (https://github.com/wwood/CoverM) with the *contig* mode, the *mean* method, and considering 95% as the minimum threshold for both sequence identity and the length of the read alignment to the lytic Microcoleus phage genomes to calculate the sequencing coverage (reads/nucleotide) in all the metagenomic samples from the Wolastoq and Eel rivers.

## Results

### Identification of phage sequences in Microcoleus-dominated mats

We found 11 and 145 assembled viral sequences with genome completeness > 50% according to *CheckV* (at least medium-quality) from the Wolastoq and the Eel River mat samples, respectively (Table S2). The lengths of the predicted viral sequences from the Wolastoq ranged from 26 to 287 Kbp, while those from the Eel River ranged between 20 and 305 Kbp. One of the viral sequences from the Wolastoq was predicted as a provirus (i.e., the viral region is flanked by host boundaries at both 5’ and 3’ ends), while the same condition was identified for 18 viral sequences from Eel River. Details of the predicted viral sequences and the estimated assembly quality are provided in Table S2.

Viral sequences with predicted temperate or virulent replication strategies were predicted in both the Wolastoq (64 and 36%, respectively) and the Eel River (70 and 26%) mats (Table S2). A low number of viral sequences from Eel River (4%) were predicted to have a chronic lifecycle (Table S2). Most of the viral sequences recovered from both rivers were classified in the *Myoviridae*, *Podoviridae* and *Siphoviridae* families (Figure 1 and Table S2). The diversity of the viral families is different in mats of each river. In samples from the Wolastoq, 55% of the viral sequences were assigned to the *Myoviridae*, while in the case of the Eel River members of the *Siphoviridae* were the most abundant (Figure 1).

**Figure 1.**
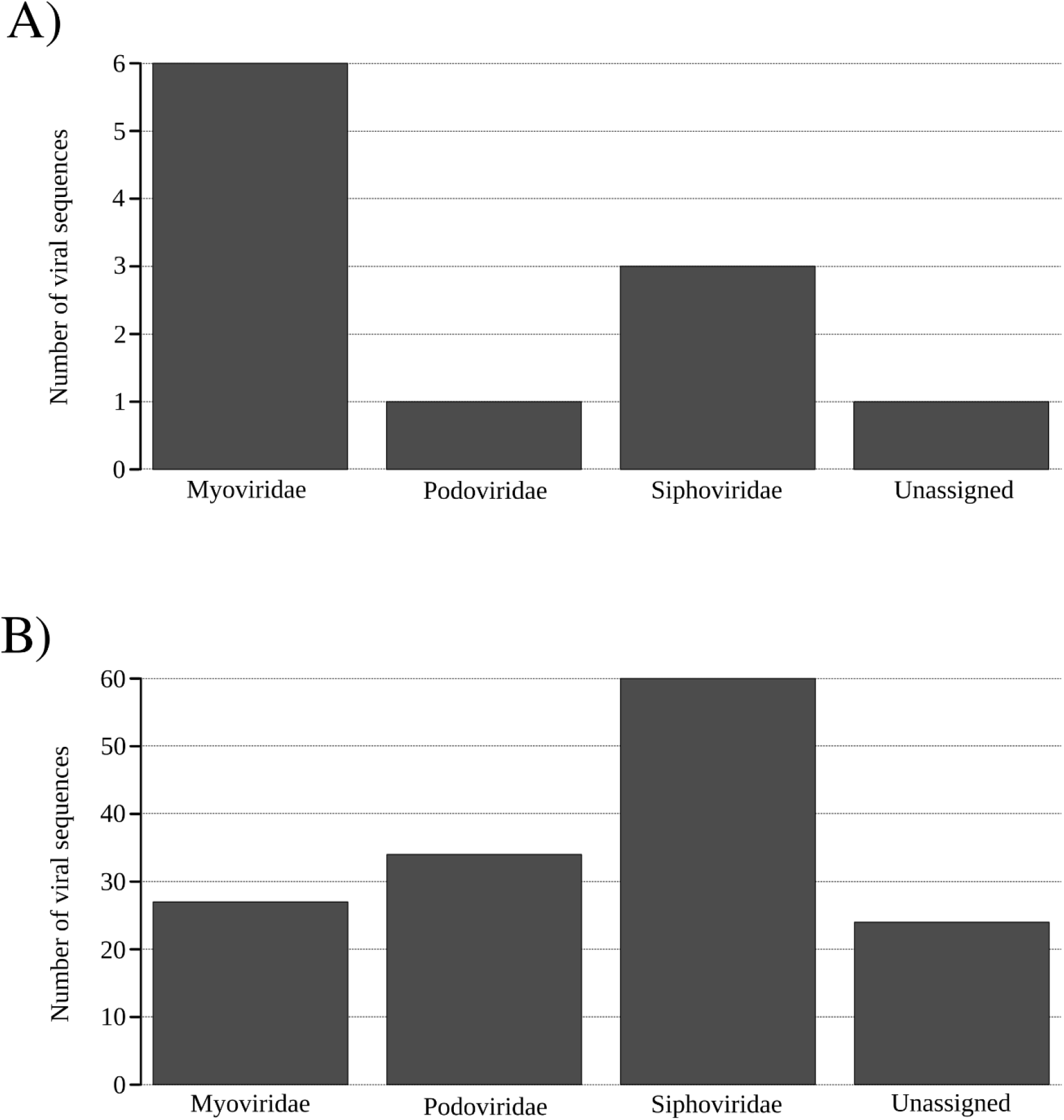
Number of assembled viral sequences from (A) the Wolastoq and (B) Eel rivers assigned to different families.

No hosts were assigned to viral sequences with the tRNA matching approach. However, the CRISPR-Cas based approach identified 11 viral sequences from the Wolastoq (3 sequences) and the Eel River (8 sequences) matching with putative bacterial hosts.

The CRISPR-Cas analysis associated phage sequences from the Wolastoq (W3-ES1) and Eel River (13US1) (both predicted as virulent phages; Table S2) with *Microcoleus* genomes (both anatoxin-producing and non-toxigenic) from New Zealand, California and Canada (Table S3). The 13US1 sequence (245,106 bp) from the Eel River was predicted as a fully assembled circular phage genome, whereas W3-ES1 (162,978 bp) appeared to be a fragment, and was recovered from the same sample as sequences W3-ES2 (30,482 bp) and W3-ES3 (45,803 bp) from the Wolastoq. Mapping W3-ES1-3 fragments against the 13US1 genome from Eel River strongly suggests that they derive from a single circa 230 Kbp phage genome. We also found spacers from *Microcoleus* genomes matching W3-ES2 and W3-ES3 sequences (Table S3), a result that reinforces the association of the predicted phage with that cyanobacteria genus.

Five prophages (06SS241, 07DS152, 07US46, 08DS177 and 08US67; predicted to have a temperate replication strategy) from Eel River (sites 6 to 8; Bouma-Gregson et al., 2019), were found integrated in assembled chromosomes of the predicted host *Microcoleus*. Other predicted phages were associated with bacterial hosts of the Proteobacteria (*Hafnia*, *Pseudomonas*, and *Klebsiella* genera), Bacteroidota, Actinomycetota and Verrucomicrobiota phyla (Table S3).

### Cyanophages residing in benthic mats of the Wolastoq and the Eel River

A phylogenetic reconstruction based on the overall similarity of protein-coding regions (“Phage Proteomic Tree”; Rohwer and Edwards, 2002) implemented in *ViPTree,* resolved the *Microcoleus*-associated phages from the Wolastoq and Eel River in two different groups predicted to infect different cyanobacterial hosts (Figure 2). In the first group, phages W3-ES1-3 (Wolastoq) and 13US1 (Eel River) were recovered as closely related sister lineages and branched with the Microcystis phages Ma-LMM01 and MaMV-DC (Figure 2). These novel phages were assigned to the same genus of the *Myoviridae* family, but are not closely related to any other known genera of the same family.

**Figure 2.**
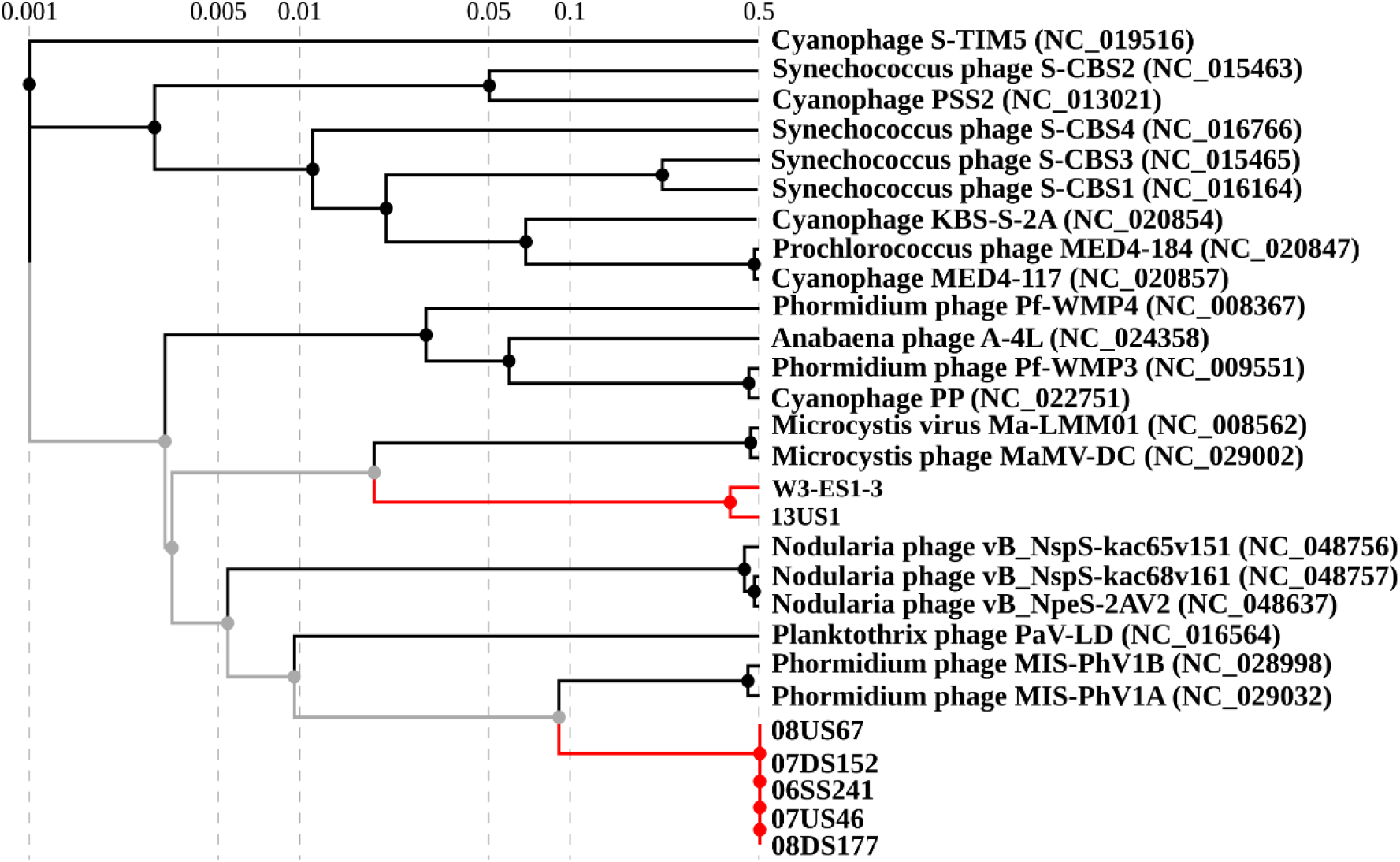
Phylogenetic relationships of diverse cyanophage genomes based on overall genomic distances. The analysis contains *Microcoleus*-associated phage genome sequences from the Wolastoq, Eel River, and diverse cyanobacteria dsDNA phages sequences (*Virus-Host DB*) as reference. The distance tree was estimated with BIONJ from a genomic distance matrix calculated with a normalized tBLASTx scoring strategy implemented in *ViPTree* (see Nishimura et al., 2017 and the Methods section). Microcoleus phages are indicated with red branches and nodes. Branch lengths are logarithmically scaled from the root (mid-point rooting). Numbers at the top indicate the log-scaled branch lengths based on the estimated genomic distance scores (Nishimura et al., 2017).

The five assembled sequences containing predicted prophages from the Eel River (Bouma-Gregson et al., 2019) have identical size (62,411 bp in all cases) and in our phylogenetic reconstruction branch with the Phormidium phages MIS-PhV1A and MIS-PhV1B (Figure 2), which were recovered from cyanobacterial mats of the Middle Island Sinkhole (Lake Huron, USA; Voorhies et al., 2016). These five new prophages were assigned to the *Siphoviridae* family and to the same genus as the Phormidium phages.

### Characteristics of the Microcoleus phage genomes

The scores defined by the Minimum Information about an Uncultivated Virus Genome standards (MIUViG) (Roux et al., 2019) classifies the *Microcoleus*-associated phages (both virulent and temperate) we report here as “high-quality draft genomes” (Table S2). These Microcoleus phage genomes lack typical plasmid signatures; therefore, it is unlikely our phage sequences represent circular extrachromosomal elements. Hereafter, we refer to the lytic phages of the *Myoviridae* family recovered from the Wolastoq and the Eel River as Microcoleus phage My-WqHQDG and Microcoleus phage My-ElHQDG, respectively. Then, the five prophages of the *Siphoviridae* family, recovered exclusively from the Eel River, are collectively referred to as Microcoleus prophage Si-ElHQDG.

The pairwise alignment of My-WqHQDG and My-ElHQDG sequences recovered from the Wolastoq (W3-ES1, W3-ES2 and W3-ES3) and the Eel River indicates that these phage genomes share multiple homologous protein-coding genes with a high sequence identity and conserved synteny (Figure 3A).

The G+C content of Microcoleus phages My-WqHQDG and My-ElHQDG is almost identical (Table 2). More protein-coding genes were predicted in Microcoleus phage My-ElHQDG (268 genes) when compared with My-WqHQDG (255 genes). Approximately 70% of the predicted ORFs are conserved between both lytic Microcoleus phage genomes (Table 2), including a set of 73 putative proteins annotated with different tools. Some of these shared protein-coding genes are highly conserved sequences (nucleotide identity > 98.5%) encoding phage structural proteins (e.g., phage major capsid, tail sheath protein and baseplate protein) and enzymes required for lysis of the host peptidoglycan cell-wall (e.g., lysin). Also, genes encoding enzymes involved in nucleotide metabolism (i.e., ribonucleotide reductase, dUTP pyrophosphatase, thymidylate synthase) and DNA replication, repair and recombination (e.g., DNA polymerase, uracil-DNA glycosylase, ATP-dependent DNA helicase and DNA primase) are common to both cyanophages (Figure 3A). Details of the Microcoleus phage gene annotation are provided in Table S4.

**Table 2.** General characteristics of lytic Microcoleus phage genomes My-WqHQDG and My-ElHQDG.

| Sampling source | My-WqHQDG | My-ElHQDG |
| --- | --- | --- |
|  | Wolastoq (NB, Canada) | Eel River (CA, USA) |
| Genome size (bp) | 239,263 | 245,106 |
| GC content (%) | 47.7 | 47.9 |
| Predicted genes | 255 | 268 |
| Shared genes (%) | 67.91 | 71.37 |
| Unique <u>genes</u> (%) | 32.09 | 28.63 |
| Genes with top viral BLAST hit | 34 | 33 |
| Annotated genes (VOG database) | 46 | 43 |

**Figure 3.**
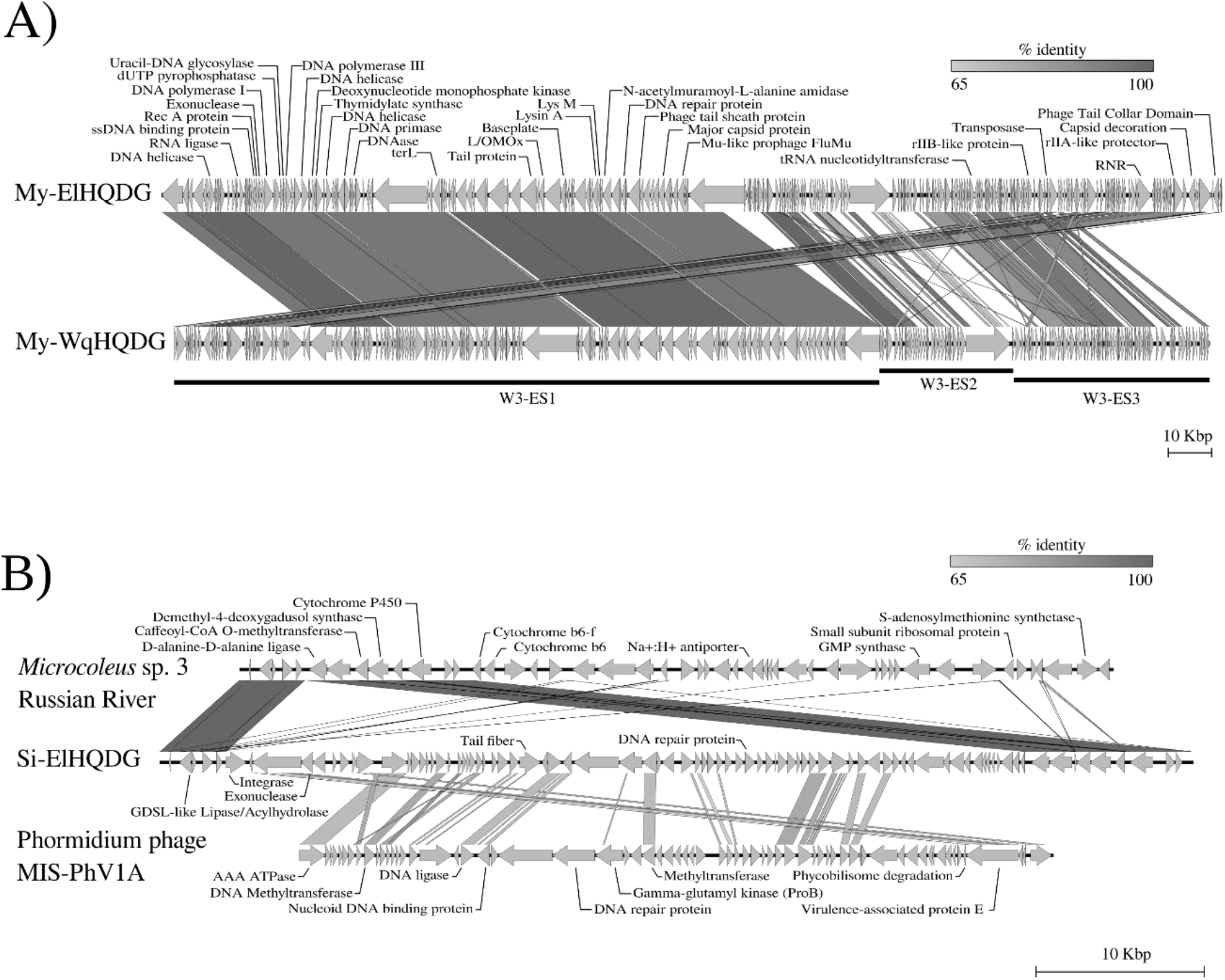
Alignments of *Microcoleus*-associated phage genomes from the Wolastoq and Eel River. **A**) Pairwise alignment of the lytic Microcoleus phages from the Wolastoq (My-WqHQDG; three fragments) and the Eel River (My-ElHQDG). Black bars in the My-WqHQDG sequences indicate the three genome fragments. **B**) Pairwise alignment of prophage Si-ElHQDG (06SS241 as reference of the identical predicted temperate phage from the Eel River) and Phormidium phage MIS-PhV1A (Voorhies et al., 2016) complete genomes and a chromosome fragment of the cyanobacteria *Microcoleus* sp. 3 from the Russian River (Bouma-Gregson et al., 2022). Grey arrowheads represent all detected ORFs, including those with no predicted annotations. Diagonal lines between genome maps represent the ORF nucleotide sequence identity (tBLASTn percent) indicated by the scale bar (% identity). Abbreviations: terL, Terminase large subunit; L/OMox, lysine/ornithine N-monooxygenase; RNR, Ribonucleotide reductase; DNAbp, DNA-binding protein.

The five predicted Si-ElHQDG Microcoleus prophages contain 86 protein-coding genes. Circa 50 Kbp of the Si-ElHQDG genome (70% of the total length) aligns with sections the Phormidium phage MIS-PhV1A (45,532 bp; Figure 3B). The Microcoleus prophages contain 30 genes homologous to coding regions found in Phormidium phage MIS-PhV1A, including sequences encoding a methyltransferase, AAA ATPase, DNA ligase, nucleoid DNA binding protein, virulence associated proteins E and ProB and the NblA protein, which is involved in the proteolytic degradation of cyanobacteria phycobilisomes during nitrogen starvation (Collier and Grossman, 1994; Yamanaka and Glazer, 1980) (Figure 3B and Table S4).

Neither end of the Si-ElHQDG prophage genome has nucleotide sequence similarity with the Phormidium phage MIS-PhV1A, but some of the contained genes are homologous with coding sequences present in a continuous segment of the chromosome of *Microcoleus* sp. 3 from the Russian River (Figure 3B).

### Microcoleus phages My-WqHQDG and My-ElHQDG are associated with toxigenic benthic mats

Except for mats W2-A and W8-A, the microbial communities of most Wolastoq samples were dominated by cyanobacteria (Figure 4A). Recently, we reported that subspecies *Microcoleus* sp. Wq-I (non-toxigenic) and *Microcoleus* sp. Wq-II (anatoxin producer), which are identical at the level of the *rps3* gene sequence, coexisted in Wolastoq benthic mats at similar relative abundances during 2018 (Valadez-Cano et al., 2023), but in the 2019 Wolastoq samples used in the current study, we only identified the toxin-producing *Microcoleus* subspecies.

**Figure 4.**
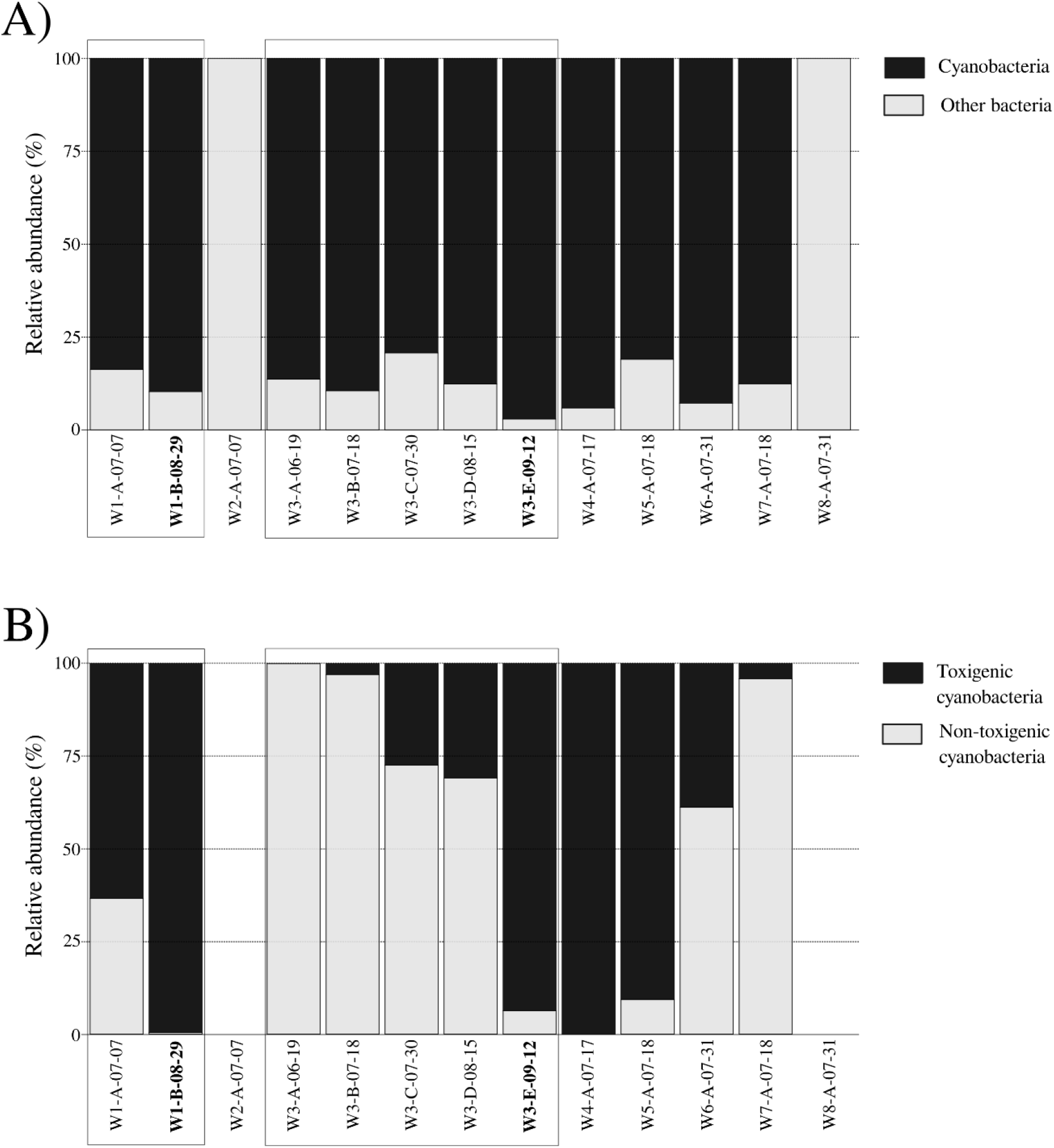
Estimated relative abundance of bacteria and cyanobacteria in Wolastoq benthic mats from 2019. **A**) Relative abundance of cyanobacteria and other bacterial lineages based on the sequencing coverage (reads/nucleotide) of the different *rps3* genes identified in benthic mats. **B**) Relative abundance of *Microcoleus* sp. Wq-II (Valadez-Cano et al., 2023) and other non-toxigenic cyanobacteria also estimated with sequencing coverage data (reads/nucleotide) of the cyanobacterial *rps3* genes identified in mat samples. Sample collection days are indicated in the four last digits of the mat names (MM-DD). Sampling sites on the Wolastoq are indicated with the W1 to W8 prefixes (same site samples are grouped in boxes). Samples where Microcoleus phage My-WqHQDG was identified are indicated in bold.

The estimated relative abundance of toxigenic *Microcoleus* sp. Wq-II varied in comparison with other non-toxigenic cyanobacterial species at different sites on the same date, and on different dates at the same site (Figure 4B). At sites 1 and 3 there was a temporal shift from dominance by non-toxigenic to toxigenic cyanobacteria as the summer progressed (Figure 4B). The detection of Microcoleus phage My-WqHQDG coincided with a very high relative abundance (94%) of toxigenic *Microcoleus* sp. Wq-II at site 3 in mid-September (cyanophage sequencing coverage 126 reads/nucleotide) (Figure 4B). Also, in mat sample W1-B, where *Microcoleus* sp. Wq-II was the dominant cyanobacterium (Figure 4B), we found reads mapping to 80% of the My-WqHQDG genome, although at a lower sequencing coverage (1.25 reads/nucleotide).

Microcoleus phage My-ElHQDG was recovered from mat sample PH2015_13U (collected upstream of site 13) from the Eel River, and we also found evidence of this cyanophage in a sample collected downstream of the same site (PH2015_13D), but with lower sequencing coverage (17 reads/nucleotide). Anatoxin-producing *Microcoleus* sp. 2 dominated the mat samples collected from that site (Bouma-Gregson et al., 2019). We did not find evidence of the lytic Microcoleus phage in scaffolds from Eel and Russian river mats collected in 2017 where only non-toxigenic species *Microcoleus* sp. 1 and *Microcoleus* sp. 3 prevailed (Bouma-Gregson et al., 2022).

### Microcoleus prophage Si-ElHQDG is integrated in mat-forming Microcoleus sp. 3 genomes from the Eel River

Microcoleus prophage Si-ElHQDG was identified in 5 samples (PH2015_06S, PH2015_07D, PH2015_07U, PH2015_08D and PH2015_08U) collected from three sites at South Fork Eel River. In all these samples we also detected the non-toxigenic *Microcoleus* sp. 3, but the toxigenic *Microcoleus* sp. was present only in three of the sampled sites (Bouma-Gregson et al., 2019).

We also explored the presence of Microcoleus prophage Si-ElHQDG in sequences assembled from Eel and Russian River mats collected in 2017 (Bouma-Gregson et al., 2022). We found the prophage in four mats collected in the South Fork Eel River, where *Microcoleus* sp. 3 was recovered. Despite finding the regions flanking the Microcoleus prophage Si-ElHQDG in *Microcoleus* sp. 3 genomes from the Russian River, we did not find evidence of the phage integrated in the cyanobacterial chromosome. Likewise, we did not detect the Microcoleus prophage Si-ElHQDG in *Microcoleus* genomes from Atlantic Canada nor New Zealand.

## Discussion

### The diverse phages inhabiting riverine benthic mats

Our metagenomic survey of cyanobacteria-dominated benthic mats from two distant North American rivers (i.e., 4550 km apart) detected 156 phage sequences from three different families and with dissimilar replication strategies. However, most of the identified phages were not linked with specific bacterial hosts. The association of phages identified in metagenome data with their specific hosts is a well-documented limitation of cultivation-independent approaches (Coclet and Roux, 2021), although computational methods based on the identification of sequence homology, such as CRISPR-spacer sequence matching, are effective at predicting phage-host relationships (Edwards et al., 2015).

We recovered cyanophages predicted to infect *Microcoleus* species from the Wolastoq and Eel River. Additionally, we identified other phages likely infecting heterotrophic bacteria that are also abundant in benthic mats. Non-cyanobacterial species that coexist with cyanobacteria in mats are predicted to play important roles in nutrient exchange and community dynamics (Bouma-Gregson et al., 2019). The heterotrophic bacteria predicted to be infected by the identified phages include members of Bacteroidetes and Verrucomicrobia, which commonly co-occur in *Microcoleus*-dominated mats from the Eel River, and proteobacteria that contribute sulfur metabolic potential to the microbial community (Bouma-Gregson et al., 2019). These findings suggest that diverse phages play important roles in shaping the microbial diversity of mats beyond the dominant cyanobacteria.

### Prophages associated with non-toxigenic Microcoleus mats from Eel River

Our computational analysis suggests that Microcoleus prophage Si-ElHQDG should be considered a member of the same genus that includes the Phormidium phages MIS-PhV1A and MIS-PhV1B, which infect *Phormidium* that dominate mats from the Lake Huron in North America (Voorhies et al., 2016). In contrast with the Microcoleus prophage, there is no evidence that the Phormidium phage sequences were integrated as prophages in the host chromosome (Voorhies et al., 2016). Prophages have been found in nearly half of the bacteria cultivated to date, particularly in pathogens and species with short doubling times, and their prevalence correlates with the genome size of the host (Touchon et al., 2016). Depending on the metabolic state of the bacterial host, temperate phage may initiate a lytic cycle or integrate into the bacterial chromosome (i.e., become a prophage) and replicate with the host chromosome (Fortier and Sekulovic, 2013). Then, under unfavorable conditions (i.e., nutritional, UV radiations, and heat shock stress), prophages may excise from the bacterial chromosome and revert to their lytic cycle (Fortier and Sekulovic, 2013). The presence of the prophage in the *Microcoleus* sp. 3 from Eel River may therefore suggest conditions in the river at the time of sampling were rather favorable.

Microcoleus prophage Si-ElHQDG was found integrated in the chromosome of *Microcoleus* sp. 3 recovered from a spatially restricted area of the Eel River in 2015 and 2017 (Bouma-Gregson et al., 2022, 2019) but not in *Microcoleus* sp. 3 from the Russian River stream, which is connected to the Eel River. Bouma-Gregson and collaborators (Bouma-Gregson et al., 2022) propose that the prevalence of *Microcoleus* sp. 3 in both rivers may be attributed to the dispersal of this species across the Eel River watershed, or it may have been present in a single ancestral river prior its division by tectonic activity about 2 million years ago (Lock et al., 2006). Restriction of the prophage to Eel River could indicate that phage dispersal is limited. However, this seems unlikely because *Microcoleus* subpopulations do not appear to be isolated in the watershed (Bouma-Gregson et al., 2022). Alternatively, temperate phage may be dispersed throughout the watershed, and the presence of prophage in Eel River genomes and absence in Russian River genomes may be due to local factors regulating the integration and/or induction of the prophage. In this way, the differential presence of prophage Si-ElHQDG may provide insight into different conditions faced by *Microcoleus* sp. 3 in each environment.

The presence of prophage can impact host fitness. Prophages can protect against superinfection by homologous phages through prophage-encoded proteins that repress phage and enhance host cell viability (Bondy-Denomy and Davidson, 2014). Similarly, prophages can prevent host superinfection by exclusion, which involves alterations of the host cell surface or other components of the cell envelope (Bondy-Denomy and Davidson, 2014). Prophages are capable of transferring genes horizontally between different bacterial hosts, such as the classic example of acquiring antibiotic-resistance (Wendling et al., 2021). Prophage integration can also cause host gene disruption, chromosome rearrangement and the expression of new functions (Fortier and Sekulovic, 2013). *Microcoleus* sp. 3 may therefore have different fitness states in different areas of the watershed due to the presence or absence of prophage.

### Lytic Microcoleus phages in toxigenic mat samples from two distant rivers

Our CRISPR-Cas system survey revealed past interactions of lytic phages My-WqHQDG and My-ElHQDG with toxigenic (i.e., the anatoxin biosynthesis gene cluster is present) and non-toxigenic *Microcoleus* species (Table S3). This is consistent with the wide host-ranges reported for some phages that can infect multiple bacterial species of the same genus, or even members of different genera (Ross et al., 2016). Given that the *Microcoleus* genomes predicted as hosts also match MAGs recovered from congeneric species from New Zealand rivers (Tee et al., 2021, 2020) (Table S3), it is feasible that the cyanophages we have identified in two riverine systems of North America are also capable of infecting mat-forming *Microcoleus* from New Zealand streams.

We did not identify toxigenic *Microcoleus* genomes with CRISPR spacers that matched cyanophage sequences recovered from the same river. For example, the only *Microcoleus* genomes with CRISPR spacers matching My-WqHQDG (Wolastoq) were the non-toxigenic *Microcoleus* sp. Wq-I from the same river, as well as toxigenic and non-toxigenic species from USA and New Zealand (Table S3). Similarly, most *Microcoleus* genomes with CRISPR spacers matching the My-ElHQDG (Eel River) were from New Zealand. This suggests toxigenic cyanobacteria in the Wolastoq and Eel River are susceptible to My-WqHQDG or My-ElHQDG infections, respectively, and their abundance influenced by infection dynamics, as has been demonstrated for *Phormidium* and phage not represented in their CRISPR defense system (Voorhies et al., 2016). Although bacteria have other strategies to prevent phage infections such as surface mutations or modifications to prevent adsorption, and restriction-modification systems to recognize and degrade phage DNA after injection (Hampton et al., 2020), the presence of My-WqHQDG and My-ElHQDG spacers in *Microcoleus* genomes from New Zealand and *M. anatoxicus* suggests that CRISPR-based protection against phage infection is important, and that these hosts have interacted in the past with viruses currently found in distant rivers. To our surprise then, we found CRISPR spacers matching the My-ElHQDG phage in the toxigenic *Microcoleus* from the Russian River, but not in the toxigenic *Microcoleus* from Eel River. This suggests the Russian River population been exposed to the lytic phage in the past and is now protected, while the Eel River population is still susceptible.

### Kill-the-winner dynamics in Microcoleus-dominated mats from the Wolastoq

The shift in dominance from non-toxigenic to toxigenic *Microcoleus* at site 3 in the Wolastoq as the summer progressed suggests that particular conditions favored the proliferation of the anatoxin-producing species and consequently of Microcoleus phage My-WqHQDG. The diversity of coexisting non-toxigenic *Microcoleus* in benthic mats, predominance of the toxigenic species or strains under specific conditions, and proliferation of the infecting phage is consistent with the “kill-the-winner” (*ktw*) dynamic equilibrium between phages and their hosts. The *ktw* hypothesis proposes a density-dependent interaction where an increase in the bacterial host population will be followed by a concomitant increase of infecting phages, which leads to a decrease in competitive hosts. This prevents a winner from emerging and ensures coexistence of competing bacteria and therefore bacterial diversity (Marantos et al., 2022; Rodriguez-Valera et al., 2009; Thingstad, 2000).

The identification of the My-ElHQDG cyanophage in Eel River mats (downstream and upstream of Site 3) dominated by toxigenic *Microcoleus* (Bouma-Gregson et al., 2019), and the absence of cyanophage sequences in all mats dominated by non-toxigenic cyanobacteria, suggests that the Microcoleus cyanophage we have identified is interacting mainly, and likely exclusively, with anatoxin-producing *Microcoleus* species of the Wolastoq and Eel River.

### Novel lytic Microcoleus jumbo phages

The large size (> 200 Kbp) of the Microcoleus My-WqHQDG and My-ElHQDG genomes places them in the range of “jumbo” phages (Yuan and Gao, 2017). Sequencing of environmental DNA from diverse geographical regions has revealed that phages with genomes > 200 Kbp infect a wide range of bacterial hosts across Earth’s ecosystems, including two jumbo phage genomes identified in mat samples from the Eel River (Al-Shayeb et al., 2020). In contrast with the abundant description of relatively small phage genomes, reports of jumbo phages are rare because their large virions commonly interfere with isolation procedures and prevent efficient separation from the host DNA using filtration methods (Yuan and Gao, 2017). Large phages have been more frequently isolated from aquatic environments that may enhance particle diffusion and therefore their ability to interact with host bacteria (Yuan and Gao, 2017).

To our knowledge, My-WqHQDG and My-ElHQDG are the first jumbo cyanophages predicted to infect *Microcoleus* hosts. Additionally, most of the jumbo cyanophage genomes reported in the *NCBI virus database* (reviewed on March 6, 2023), the majority infecting *Prochlorococcus* and *Synechococcus*, were recovered from marine environments. Some of these large phages prey on freshwater cyanobacteria associated with bloom formation and toxin production. For example, the jumbo cyanophage PhiMa05 (273.8 Kbp) is highly efficient at infecting microcystin-producing *Microcystis* isolates from lakes (Naknaen et al., 2021). This PhiMa05 cyanophage is also classified in the *Myoviridae* family (Naknaen et al., 2021), but is not related to My-WqHQDG and My-ElHQDG.

The average size of the Microcoleus My-WqHQDG and My-ElHQDG genomes is approximately 70 Kbp longer than cyanophage MaMV-DC (∼169.2 Kbp), which infects *Microcystis* species (Ou et al., 2013; Wang et al., 2019). It has been proposed that jumbo phages evolved from smaller phage ancestors through the recruitment of novel genetic information (i.e., nonhomologous recombination between unrelated sequences and recruitment of host genes by horizontal gene transfer) that has increased both the genome size and genetic capabilities (Hendrix, 2009). Larger virions and greater coding capacity typically results in more genes substituting the functions of host proteins, thus reducing the dependency on their hosts and thereby increasing host range (Yuan and Gao, 2017). In contrast with other cyanophages, we did not find evidence of genes associated with host functions such as photosynthesis (i.e., PSI and PSII components), phycobilisome degradation (i.e., nblA), carbon metabolism, or phosphorus-acquisition (Gao et al., 2016).

### Phages and their role in riverine benthic microbial communities

Phages can be important drivers of diversity and abundance in microbial communities. For instance, it has been estimated that phages lyse 20-40% of the bacterial cells in marine surface water on a daily basis (Suttle, 2007, 1994). Our results suggest that anatoxin-producing *Microcoleus* are especially susceptible to phage infection. Microcoleus My-WqHQDG and My-ElHQDG phages are predicted to have a lytic replication cycle, suggesting that they can influence the proliferation and abundance of *Microcoleus* species that dominate benthic mats and drive toxin production potential.

Phage-induced lysis releases intracellular products, strongly influencing the microbial food web and biogeochemical cycling (Weinbauer, 2004). Anatoxins are consistently released from *M. autumnalis* into the water (Wood et al., 2018). Factors associated with cyanotoxin release include substrate/metabolite active transport, as suggested for microcystin-producing *Microcystis aeruginosa* PCC 7806 (Pearson et al., 2004), and cell lysis due to physicochemical factors within mats for anatoxin-producing *Microcoleus* (Wood et al., 2018). Infection of microcystin-producing *Microcystis* with the jumbo cyanophage PhiMa05 resulted in release of microcystins as consequence of lysis, and microcystin production (intracellular and extracellular) was reduced when compared to uninfected cells (Naknaen et al., 2021). Likewise, lytic Microcoleus phages may play an important role in the production of anatoxins by *Microcoleus* and their release into the surrounding environment.

In addition to the Microcoleus phages, we found other phages that could potentially infect other bacteria associated with the benthic mats. In aquatic systems, a significant fraction of the prokaryotic community is infected with phages (Weinbauer, 2004), a process that contributes to microbial mortality rates, reshapes bacterial communities and influences the long-term evolution of bacterial populations (Coclet and Roux, 2021). Microbial assemblages in *Microcoleus*-dominated mats consist of multiple bacteria co-occurring with cyanobacteria (Bouma-Gregson et al., 2019). Hence, phages infecting non-cyanobacterial, but abundant, hosts may also have an important role in the microbial community dynamics and proliferation of toxigenic or non-toxigenic mat-forming cyanobacteria.

## Conclusions

To our knowledge, this is the first description of cyanophages associated with toxigenic mat-forming *Microcoleus* species from freshwater environments. The discovery of these phages highlights the relatively unexplored microbial communities of freshwater systems, in particular benthic environments, and the key roles played by phage therein. Cyanophage infections may influence the population dynamics of anatoxin-producing cyanobacteria that dominate benthic mats in rivers around our planet and pose a threat to wildlife, companion animals, and humans that recreate in these waters.

## Supplementary figures

**Figure S1.**
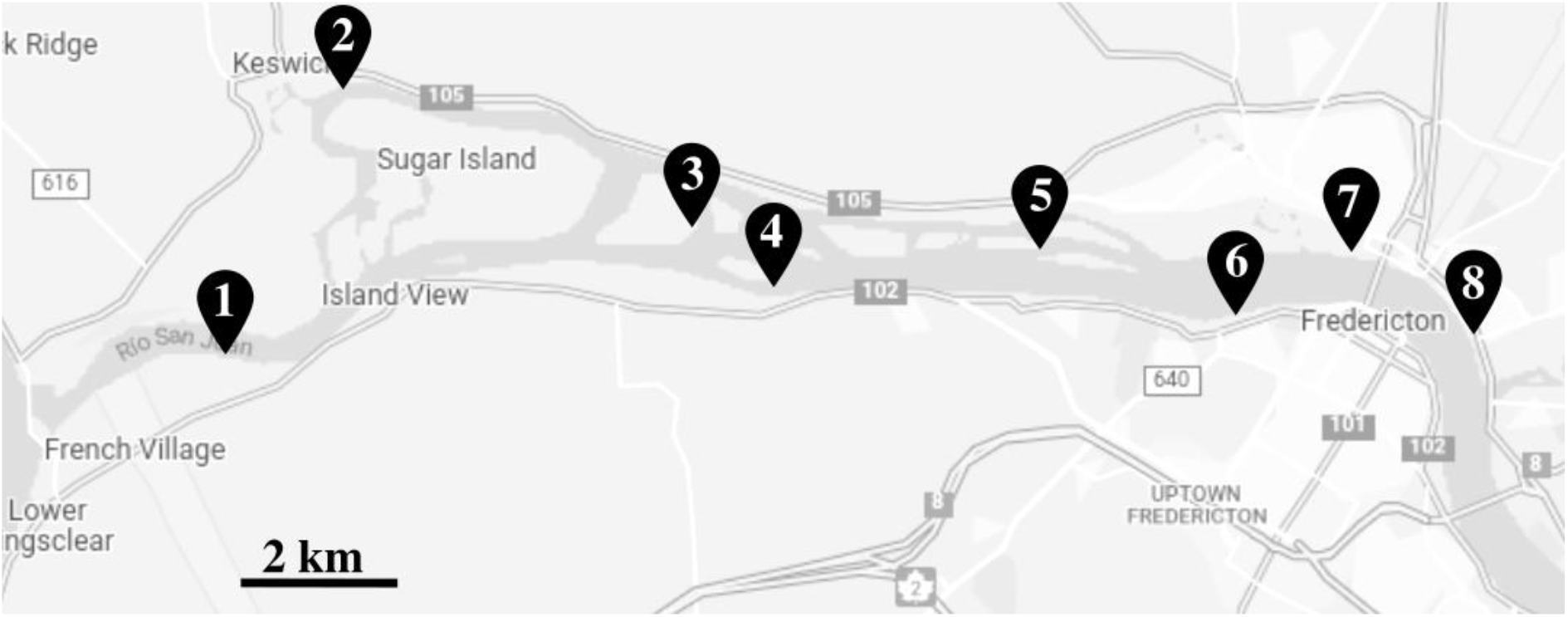
Map illustrating the 2019 sampling sites in the Fredericton region (New Brunswick; Canada) of the Wolastoq (Saint John River).

## Supplementary table legends

Table S1. Metagenome-assembled genomes analyzed.

Table S2. Viral sequences recovered from metagenomic data from the Wolastoq and Eel rivers.

Table S3. CRISPR spacers of different bacterial genomes matching viral sequences recovered from the Wolastoq and Eel rivers. Genome names of toxigenic Microcoleus are indicated in bold.

Table S4. Details of annotated genes in the lytic and lysogenic Microcoleus phage genomes.

